# Phosphorus stress and spatial confinement lower the threshold for quorum-sensing activation of redox-active metabolite production in *Pseudomonas synxantha*

**DOI:** 10.1101/2025.11.11.687776

**Authors:** R. E. Alcalde, H. Jeckel, O. Zhang, R. L. Avila, N. Dalleska, D. Mavrodi, C. Yang, D. K. Newman

**Affiliations:** Division of Biology and Biological Engineering, California Institute of Technology, Pasadena, CA 91125; Department of Physics & Astronomy, Texas A&M University, College Station, TX 77843; Global Environmental Center, California Institute of Technology, Pasadena, CA 91125; USDA, Agricultural Research Service, Wheat Health, Genetics and Quality Research Unit, Pullman, Washington, USA; Division of Engineering and Applied Science, California Institute of Technology, Pasadena, CA 91125; Division of Geological and Planetary Sciences, California Institute of Technology, Pasadena, CA 91125

## Abstract

Bacteria often coordinate collective behaviors such as biofilm formation and secondary metabolite production through quorum sensing (QS), a regulatory system traditionally linked to high cell density. However, in environments like soil, where microbial populations are spatially fragmented, sparse, nutrient-limited, and subject to mass transport, the mechanisms that enable QS-dependent processes remain incompletely understood. Here, we investigate the regulation of a secreted redox-active metabolite, phenazine-1-carboxylic acid (PCA), in *Pseudomonas synxantha* 2-79, a model rhizobacterium, under phosphorus (P) limitation, a persistent stress in many soils. Using a combination of microscopy and molecular genetic approaches, we show that P limitation sensitizes the QS activation threshold by an order of magnitude, enabling phenazine induction at relatively low population densities in comparison to P replete conditions. This induction is abolished in QS-deficient mutants and restored by the addition of exogenous acyl-homoserine lactone (AHL), demonstrating that quorum sensing remains essential but its threshold becomes environmentally tuned. Under P limitation, spatial confinement and pore saturation levels further shape the timing and location of induction, illustrating how physical structure and nutrient stress can modulate bacterial activities. Moreover, phosphorus stress confers both collaborative and competitive advantages, enabling *P. synxantha* to undergo low-cell density AHL cross-talk between related *Pseudomonas* spp. and to suppress other rhizobacteria. Lastly, on plant roots, PCA genes are more predominantly induced under P limitation. These findings illustrate how the nutrient status of an environment can modulate the onset of quorum sensing, enabling quorum-regulated behaviors to activate at lower thresholds.

## Introduction

Soil microbes sustain ecosystems. Through their metabolism, they drive nutrient cycling, modulate plant health, and influence mineral weathering. Many of these functions depend not only on individual physiology but on the coordinated activity of microbial populations and communities, often mediated through the release of small molecules that act as public goods, chemical signals, or antibiotics. Among these, phenazines—redox-active metabolites produced by diverse soil bacteria that support a variety of biological functions^1–4^—are particularly notable. Through their redox chemistry, they promote phosphorus^5,6^ and iron availability,^7–10^ shape microbial competition,^11–13^ and affect plant-microbe interactions^14–16^ in the rhizosphere.

Phenazine biosynthesis is typically controlled by quorum sensing (QS), a gene regulatory system in which bacteria produce and respond to diffusible signals, such as acyl-homoserine lactones (AHLs), to coordinate group-level behaviors.^17,18^ Once a threshold concentration of signal accumulates, often reflected by high cell density, QS activates gene expression for costly cooperative functions like biofilm formation, virulence, or secondary metabolite production.^19–25^ This framework is well supported in laboratory systems. However, in natural environments like soil, microbial populations are often spatially fragmented, sparse, and physically confined within pore structures,^26–28^ conditions that can shift signal accumulation and QS activation. Cluster analyses in structured soils report that larger clusters (>100 cells) become sparse as water content decreases, implying many neighborhoods fall below 100 cells.^26^ Despite this, phenazines^29,30,4^, and other QS-regulated metabolites^31–33^ are commonly detected in the rhizosphere. How, then, are these behaviors triggered in environments where signal accumulation may be restricted?

Several models emphasize that QS dynamics also depend on environmental structure and signal transport. Diffusion sensing and efficiency sensing, alternative descriptions of quorum sensing, describe how mass transfer, spatial confinement, and local cell organization determine whether signal concentrations reach activation thresholds.^34–37^ In structured habitats like biofilms and soils, spatial confinement and mass transfer limitation may allow signals to accumulate locally even when bulk population density is low.^28,38–41^ Consistent with this view, a few single cells confined in picoliter microfluidic chambers initiate “high-density” QS programs.^38^ Likewise, flow cell experiments show that advection suppresses QS in exposed regions of a biofilm while sheltered niches maintain activation.^42^

Beyond physical structure and transport, rhizosphere chemistry and biology, such as nutrient limitation, physicochemical stress, and host-derived cues can also tune QS induction^43^. Here we focus on phosphorus limitation in *Pseudomonas synxantha* (formerly *P. fluorescens*) 2-79, a plant growth-promoting rhizobacterium that produces phenazine-1-carboxylic acid (PCA).^44,45^ In this strain, PCA production is governed by a single AHL-based quorum-sensing system, PhzIR; deletion of either *phzI* or *phzR* abolishes PCA production.^45–47^ Although PCA increases under phosphorus limitation,^6,48^ whether this induction is mediated by the PhzIR circuit, arises via cross-talk with other QS pathways, or is driven independently of QS remains unknown. This organism-level question is motivated by the broader soil context: freely available inorganic phosphate is extremely low because much of it is immobilized through sorption to mineral surfaces, ^49–51^ and in *Pseudomonas aeruginosa*, phosphate (and iron) limitation is known to influence secondary metabolism through multiple QS pathways (e.g., PQS, IQS, RhlR)^52–58^ via the PhoBR two component that governs system central phosphate sensing and response.^59–61^

We set out to determine whether phosphorus stress alters QS dynamics in *P. synxantha* 2-79. Specifically, we asked whether phosphorus-limited induction of PCA is QS-dependent or QS-independent. We postulated that phosphorus stress would trigger AHL-based PhzIR circuits, thereby lowering the effective QS cell density threshold because PCA production under P limitation occurs at lower cell densities in comparison to standard conditions.^48^ We further hypothesized that structured environments act with phosphorus levels to shape the onset, location, and ecological consequences of QS-regulated behaviors. To test these predictions, we integrated genetic, optical, and ecological approaches to better understand the rules governing microbial coordination in nature.

## Results

### Phosphorus limited environments enhance phenazine gene expression of *Pseudomonas synxantha* in plant root microcosms

To begin exploring how phenazine biosynthesis responds to phosphorus availability in the rhizosphere, we imaged *Pseudomonas synxantha* 2-79 colonizing *Brachypodium distachyon* Bd21-3 roots under phosphorus-replete (P-rep; excess inorganic phosphate in medium), phosphorus-limited (P-lim; no inorganic phosphate in medium), and phosphate-sequestering (P-HFO; phosphate bound to iron, i.e., not bioavailable) conditions using light sheet fluorescence microscopy (LSFM). In many Fe-oxide–rich, acidic to neutral soils, phosphate is strongly retained by iron oxides (e.g., hydrous ferric oxide, HFO),^50,62,63^ so our P–HFO treatment approximated this sequestration. A dual fluorescent reporter was used to track phenazine biosynthesis gene expression (P*_phzA_*-GFP, green channel) and colonization density readout (constitutive, P*_tac_*-mCherry, red channel). Fluorescence intensity was quantified from maximum intensity projections (MIPs) of LSFM images. For each object within a region of interest, the mean intensity was calculated as integrated density divided by object area. This metric approximates fluorescence per unit area in 2D projections, serving as a readout for relative reporter expression and cell density in colonized regions. Bar plots and Empirical Cumulative Distribution Functions (ECDFs) revealed condition-specific differences in both fluorescence magnitude and distribution (Fig. 1A–B).

**Figure 1:**
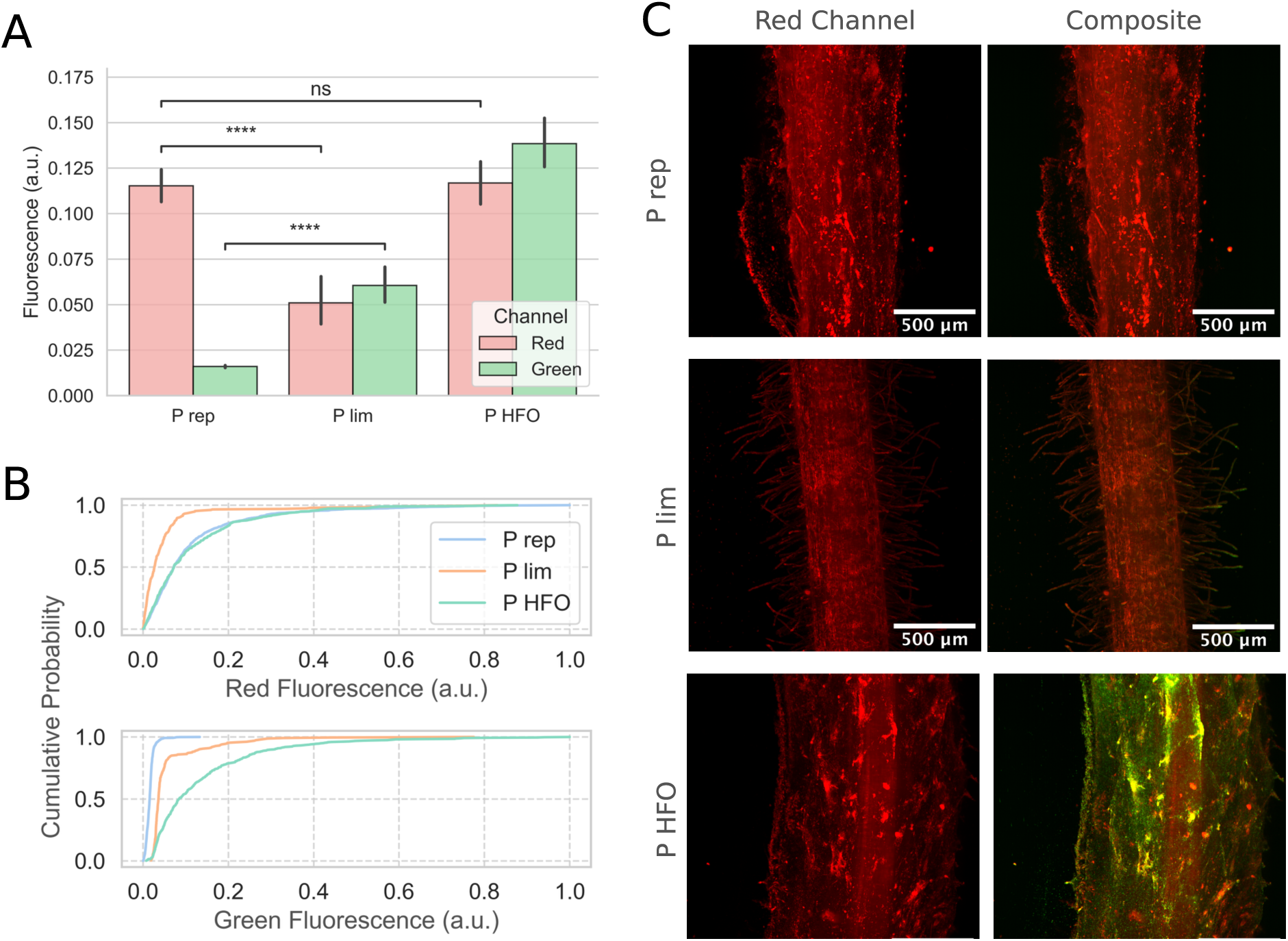
Phenazine gene expression of *Pseudomonas synxantha* 2-79 on plant roots under different phosphorus conditions. **(A)** Bar plots showing mean ± 95% CI of normalized red channel (constitutive; P*_tac_*) and green channel (phenazine biosynthesis; P*_phzA_*) fluorescence intensities under phosphorus-replete (P-rep), phosphorus-limited (P-lim), and phosphate-sorbing hydrous ferric oxide (P-HFO) conditions. Statistical comparisons were performed using two-sided Mann–Whitney tests; significance is indicated by asterisks (ns, not significant; **** p < 0.0001). **(B)** Empirical cumulative distribution functions (ECDFs) of normalized fluorescence intensities from the same dataset shown in (A), illustrating the distribution of object-level fluorescence values across experimental conditions. **(C)** Representative maximum intensity projection (MIP) images acquired by light sheet fluorescence microscopy (LSFM) of *P. synxantha* colonizing plant roots under P-rep, P-lim and P-HFO conditions. Red channel images (left) are shown with identical intensity thresholds to compare red fluorescence. Composite images (right) overlay red and green channels using the same fixed thresholds across conditions, revealing differences in P*_phzA_* reporter induction. Scale bars = 500 μm. See also Movie S1 for a 3D reconstruction, Table S1 for summary statistics, and Figure S1 for object-level strip plot.

Induction of phenazine biosynthesis (P*_phzA_*) was highest under the P-HFO condition (mean = 0.138), followed by P-lim (mean = 0.061), while remaining minimal under P-replete conditions (mean = 0.016). Although the P-lim condition exhibited lower average expression than P-HFO, its distribution was markedly right-skewed and heavy-tailed, indicating rare but strong induction events. In contrast, P-HFO displayed a broader, moderate distribution, while P-rep remained narrowly distributed around low fluorescence values (Fig. 1B). Representative LSFM images corroborated these findings, revealing robust activation of the P*_phzA_* reporter under P-HFO but not under P-replete conditions (Fig. 1C). Interestingly, phenazine expression did not directly correlate with cell density. The constitutive reporter was highest in P-HFO and P-rep (means = 0.117 and 0.115, respectively) and lower under P-lim (mean = 0.051), yet only P-lim and P-HFO conditions induced the phenazine reporter.

These observations indicate that at the time of measurement, the cell density was insufficient to trigger phenazine gene expression under nutrient-replete conditions. How, then, does phosphorus limitation stimulate phenazine gene induction at local colonization levels that fail to trigger expression under phosphorus-replete conditions? This observation suggests that canonical QS induction of phenazines may be altered under nutrient stress, but whether this results from a bypass of QS regulation, an increased sensitivity to quorum signals, or integration with parallel stress-response pathways was unclear. We set out to disentangle these possibilities.

### Quorum sensing is required for phenazine production under phosphorus limitation, and phosphorus stress sensitizes the QS system to AHL

To determine whether phenazine production under phosphorus limitation requires the *phzIR* quorum sensing (QS) pathway, we examined wild type and mutants defective in either the phenazine production genes (*phzA-G*), the QS signal synthase (*phzI*), or the phosphorus starvation response regulator (*phoB*) under P-rep (7 mM Pi) or P-lim (100 µM Pi) conditions in 96-well plate liquid cultures (Fig. 2A). The wild type produced phenazines under both conditions but showed markedly higher induction per cell under P-lim. In contrast, the Δ*phzI* mutant failed to produce phenazines under P-lim and P-rep, confirming that QS is required for induction under P-lim. Similarly, the Δ*phoB* mutant did not produce phenazines under P-lim but retained production under P-rep, indicating that phosphorus stress signaling via PhoBR is also required in the P-lim response.

**Figure 2:**
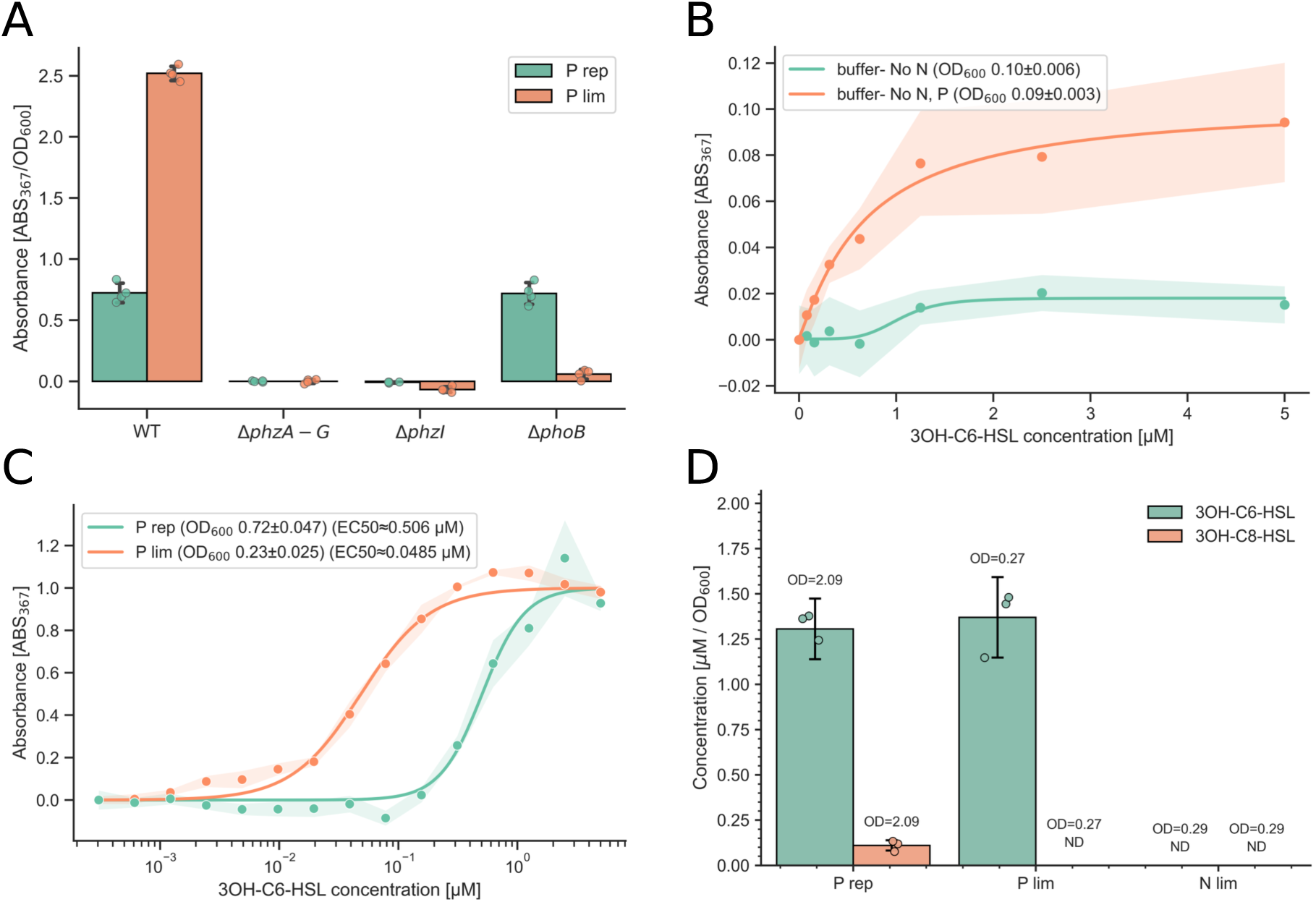
Quorum sensing requirement and reduced AHL concentration threshold for phenazine production in *Pseudomonas synxantha* 2-79 under phosphorus limitation. (**A**) Phenazine-associated absorbance normalized to cell density (ABS_367_ · OD_600_ ^-1^) in *P. synxantha* strains under phosphorus-replete (P-rep) and phosphorus-limited (P-lim) medium at 20 h post-inoculation. Values were baseline-corrected using the Δ*phzA-G* mutant mean for each condition. Bars show mean ± SD of replicates; points represent individual measurements; n=4. **(B)** Dose–response of the Δ*phzI* mutant strain to exogenous 3-OH-C6-HSL under nitrogen-free (No-N) and combined nitrogen- and phosphorus-free (No-N, P) conditions at 20 h post-inoculation. Responses were baseline-corrected by subtracting the mean signal at 0 µM 3-OH-C6-HSL within each condition. Points show per-dose means; shaded regions indicate mean ± SD of replicates; n=5. Curves show four-parameter logistic model fits. Insert in legend: mean ± SD of OD_600_ at the assay timepoint. **(C)** Dose–response of the Δ*phzI* mutant strain to exogenous 3-OH-C6-HSL at 20 h under P-rep and P-lim medium at 20 h post-inoculation. Responses were baseline-corrected and scaled to an empirical E_max_ (median of the two highest doses) to yield 0–1 normalization. Points show per-dose means; shaded bands indicate ±SD across replicates; n=3. Curves are two-parameter Hill fits; Insert in legend: mean ± SD of OD_600_ at the assay timepoint and EC_50_ values from the fits. **(D)** AHL concentration normalized to cell density (µM · OD_600_^-1^)) for 3OH-C6-HSL and 3OH-C8-HSL across P-rep, P-lim, and nitrogen-limited (N-lim) conditions. Bar plots show mean ± SD. Values above each bar indicate the corresponding optical density (OD_600_) at sampling time (22 h). “ND” denotes values below the detection limit.

We next asked whether exogenous addition of the QS signal produced by PhzI would complement Δ*phzI* and whether this response is specific to phosphorus depletion. We performed a dose–response assay by supplementing the Δ*phzI* mutant with the N-(3-hydroxy-hexanoyl)-L-homoserine lactone (3-OH-C6-HSL) (Fig. 2B). *P. synxantha* produces at least six AHLs, including three of the hydroxy form, with 3-OH-C6-HSL having the highest sensitivity to the PhzR.^46^ To restrict growth and isolate regulatory effects, we used a defined buffer medium lacking either nitrogen (No-N) or both nitrogen and phosphorus (No-N/P). After 20 h, PCA production was absent in No-N cultures, even at high QS signal concentrations, but was strongly induced in No-N/P cultures in a dose-dependent manner.

We further asked whether phosphorus limitation increases sensitivity to QS molecules. We therefore performed a second 3-OH-C6-HSL dose–response assay (Fig. 2C), this time allowing the Δ*phzI* mutant to grow under P-rep (7 mM) and P-lim (100 µM) conditions. After 20 h, PCA production was present in both conditions in a dose-dependent manner, but under P-lim, lower QS signal concentrations were sufficient to induce phenazine production. Fitting a two-parameter Hill (log-logistic) function revealed half-maximal effective concentrations (EC50) of 0.51 µM (P-rep) and 0.05 µM (P-lim), indicating ∼10-fold greater sensitivity under phosphorus stress.

Lastly, to verify QS signal production, we directly quantified AHLs (i.e., 3OH-C6-HSL, 3OH-C8-HSL, and 3OH-C10-HSL) by LC-MS under P-rep, P-lim, and N-lim (nitrogen limited) conditions for both the wild-type strain and Δ*phzI*. Results are shown for WT (Fig. 2D). For the wild-type strain, QS signals were produced under both P-lim and P-rep but were not detected under N-lim. No AHLs were detected with the Δ*phzI* strain. 3OH-C10-HSL was not detected under any conditions. Our limit of detection was 10 nM for all AHLs.

Together, these results demonstrate that phenazine production under P-lim is QS-dependent and requires the PhoBR two-component system. Moreover, phosphorus stress sensitizes QS by lowering the AHL activation threshold, enabling induction at levels that are non-inducing under P-rep or N-lim.

### Phosphorus limitation enables quorum sensing activation at low cell densities in planktonic culture

Because QS is required for phenazine production under P-lim, we inferred that phosphorus stress decreases the cell-density threshold for QS activation. To test this, we used reporter strains to track QS-dependent induction in wild type *P. synxantha* 2-79 grown in well-mixed liquid culture across phosphate regimes (25, 100, and 7000 µM Pi) with or without exogenous 3-OH-C6-HSL (AHL) (Fig. 3A-B). Our previously constructed P*_phoA_* and P*_phzA_* reporter strains^48^ served to monitor phosphorus stress (activation of PhoBR) and phenazine production, respectively.

**Figure 3:**
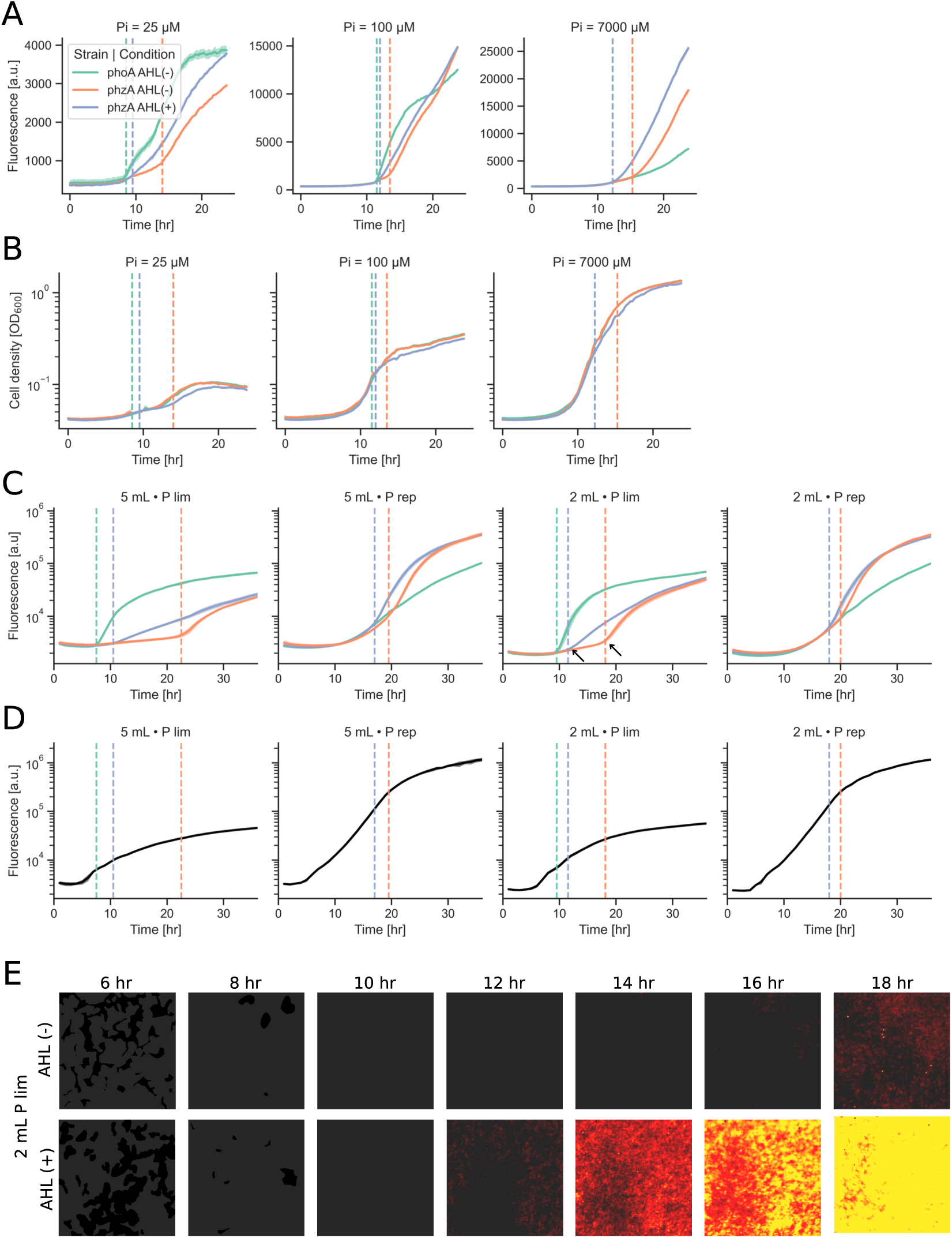
Induction benchmarks and growth dynamics under phosphorus regimes of planktonic and sessile *Pseudomonas synxantha* 2-79 cultures. (**A**) Induction curves for phosphorus-stress (*phoA*) and phenazine biosynthesis (*phzA*) reporters in liquid cultures across phosphate regimes (Pi = 25, 100, 7000 µM). Raw fluorescence intensity values are shown. **(B)** Growth curves in liquid cultures as cell density (OD_600_). **(C)** Induction curves for *phoA* and *phzA* in agar biofilms under P-lim and P-rep conditions in either 5 mL or 2 mL agar volumes. Raw fluorescence intensity values are shown. **(D)** Growth curves in agar biofilms represented by the constitutive reporter fluorescence intensity (cell density readout). **(E)** Representative images of reporter induction time course for biofilms grown under P-lim in a 2 mL agar volume. Intensity of fluorescence is depicted by red to yellow hue. Gray shading denotes the area coverage of aggregates. Legend: WT P*_phoA_* without AHL addition (green), WT P*_phzA_* without AHL addition (orange), P*_phzA_* with AHL addition (blue). Curves denote mean; shading denotes ± SD; n = 3. Error not seen is smaller than the line. Dashed vertical lines mark reporter induction times. Arrows on third graph correspond to images in (E)

The *phoA* reporter was induced in mid-exponential phase at 25 and 100 µM Pi (P-lim), with both induction timing and cell density increasing with phosphorus availability. It was not detected at 7000 µM Pi (P-rep). Without added AHL, the *phzA* reporter was induced in late exponential/early stationary phase under all conditions, but earlier and at a substantially lower cell density in P-lim compared to P-rep (OD_600_ at induction: 25 µM, 0.0846 ± 0.001; 100 µM, 0.219 ± 0.005; 7000 µM, 0.806 ± 0.006; mean ± SD), representing ∼10-fold and ∼4-fold reductions at 25 and 100 µM, respectively. Addition of exogenous AHL advanced induction time and lowered the density threshold further; under P-lim. Moreover, *phzA* onset aligned more closely with *phoA*, consistent with phosphorus stress priming the QS response while endogenous signal accumulation is rate-limiting when AHL is not supplied. It is also noteworthy that even when AHL is exogenously added in excess, gene expression still requires the cells to be under P limitation or relatively high cell density under P-rep for induction. That is, it seems that beyond requiring high AHL concentrations the cells need to be in a state of physiologically readiness (e.g., P stress, other stress responses at high cell density).

These results indicate that phosphorus stress reduces the cell-density requirement for quorum activation and speeds QS-dependent gene expression in well-mixed cultures, motivating tests in structured environments that better simulate rhizosphere matrices, where mass transfer and bacterial confinement could further tune this sensitization.

### Phosphorus limitation also lowers the cell density threshold for quorum sensing activation in sessile populations

We employed an early-stage colony–biofilm assay using our previously developed imaging system^48^ to quantify growth and reporter expression in *P. synxantha*. In this diffusion-dominated context, AHL concentrations in the biofilm arise from the balance between signal production, diffusive loss into the underlying agar (which serves as a sink), and retention within the biofilm matrix. Reporters on phosphorus stress (P*_phoA_*) and phenazine production (P*_phzA_*) tracked induction, and a constitutive reporter (P_PA10403_) provided a readout for cell density (Fig. 3C-E). By varying nutrient status, agar volume, and AHL supplementation, we modulated regulatory sensitivity, sink capacity (5 mL vs 2 mL agar volumes), and signal availability, respectively, to reveal how these factors shape QS activation.

5 mL agar: Under P-lim, *phoA* was detected early (∼7.5 hr), but *phzA* induction was markedly delayed (∼22.5 hr). Under P-rep, *phoA* did not induce (as expected), whereas *phzA* induced at 19.5 h. This pattern is opposite to planktonic cultures, where P-lim induces earlier than P-rep, indicating that in this heterogeneous, diffusion-dominated context, QS-signal accumulation is delayed under P-lim.

Notwithstanding the delay, the biofilm density required for *phzA* activation was ∼9-fold lower under P-lim than P-rep (constitutive reporter [a.u.]: P-lim 27,852 ± 1,207; P-rep 248,856 ± 13,115; mean ± SD), which is consistent with planktonic trends.

2 mL agar: Under P-lim, *phoA* was, again, detected early (∼9.5 hr), *phzA* induction occurred (∼18 hr; earlier than 5 mL condition), and before P-rep (∼20 hr), suggesting that QS-signal thresholds were achieved more rapidly when sink capacity was reduced (i.e., lower agar volume). Interestingly, the biofilm density at the time of *phzA* induction was nearly identical between the 2 mL and 5 mL conditions, for both P-lim and P-rep, indicating that biomass accumulation was faster under 2 mL, likely contributing to the earlier induction. Lastly, as in the 5 mL agar experiments, P-lim, again, required ∼10-fold lower cell density than P-rep for *phzA* activation.

The addition of exogenous AHL caused P-lim colonies to induce *phzA* expression rapidly, coincident with the onset of phosphorus stress and bypassing the need for signal accumulation through growth. AHL rescue in P-rep colonies also accelerated the induction time but still required a higher cell density than P-lim. This is again consistent with liquid cultures. Exogenous AHL accelerated induction but did not bypass the requirement for phosphorus limitation or a high biofilm density.

Together, these results show that QS activation in colony biofilms reflects an interplay between nutrient status and the balance of AHL signal accumulation versus diffusive loss in the surrounding agar.

Phosphorus limitation lowers the biofilm density required for QS activation, but induction still depends on local signal concentration, which is influenced by retention within the biofilm versus loss into the agar.

### Phosphorus stress and spatial confinement accelerate quorum sensing in soil-like microenvironments

To examine how the physical structure of the environment shapes local signal accumulation and QS activation, we used a previously designed microfluidic platform that mimics soil-like porous media^64^ (Fig. 4A). Low-density cultures (OD_600_ = 0.05) of a dual reporter strain (*P. synxantha* 2-79 P*_tac_*-P*_phzA_*) were inoculated and air was introduced to generate a landscape of water pockets that varied in size, connectivity, saturation levels (Fig. 4B). Across roughly 50 regions of interest (ROIs) imaged in each experiment, we tracked biomass (P*_tac_*) and *phz* induction (P*_phzA_*) (Fig. 4C, Fig. S2).

**Figure 4:**
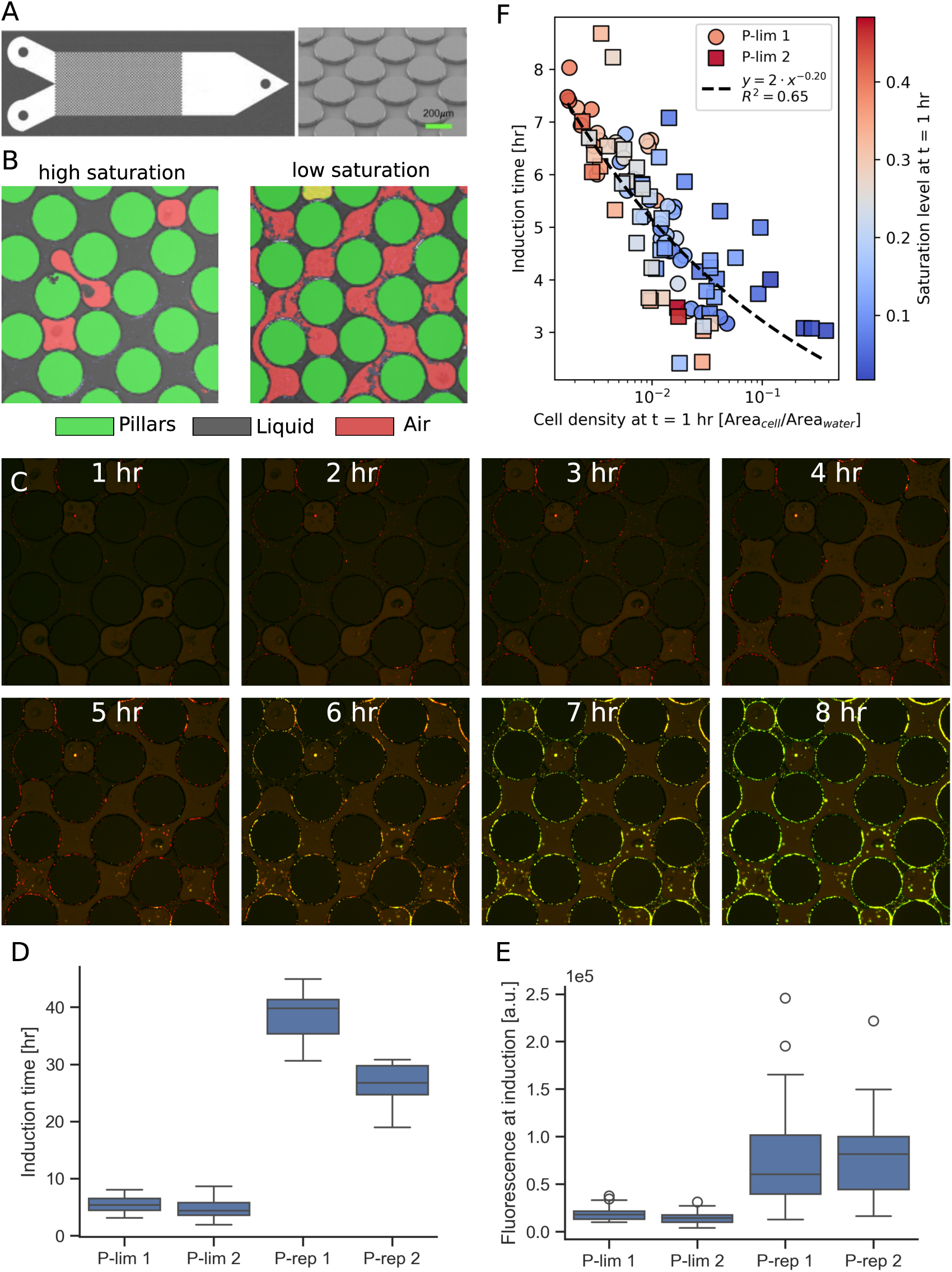
Relationship between cell density, water saturation, and *phz* induction time in microfluidic microenvironments. **(A)** Schematic (left) and scanning electron micrograph (right) of the synthetic porous platform used to create spatially structured microenvironments. The reactor consists of a 2 by 1 cm pore network containing an array of evenly spaced cylindrical posts 300 µm in diameter, a depth of 10 µm, and a porosity of 0.39. **(B)** Representative images of pore-scale water distribution at high saturation and low saturation levels, with pillars shown in green, water-filled pores in gray, and air-filled pores in red. **(C)** Representative composite images of reporter induction (constitutive: red, *phzA*: green) over time. **(D)** Boxplots summarizing *phz* reporter induction times for individual ROIs across replicate P-lim (P-lim 1–2) and P-rep (P-rep 1–2) experiments. Boxes show medians and interquartile ranges; whiskers extend to 1.5× the interquartile range. See Figure S2 for induction time estimation. **(E)** Boxplots summarizing red-channel integrated fluorescence per ROI (density proxy) at the time of *phz* reporter induction across replicate P-lim (P-lim 1–2) and P-rep (P-rep 1–2) experiments. Boxes show medians and interquartile ranges, and whiskers extend to 1.5× the interquartile range. **(F)** Distribution of induction times under P-lim. Relationship between cell density in pore water (x-axis; area fraction of cells in water-filled regions at t = 1 hr) and *phz* induction time (y-axis) for P-lim 1 (circles) and P-lim 2 (squares). Points are colored by water saturation (fraction of pore volume filled with water), as indicated by the color bar. The dashed line shows a power-law fit to all data. Data represent pooled technical replicates from regions in the reactor (P-lim 1, n = 44 ROIs; P-lim 2, n = 52 ROIs).

Comparing P-lim and P-rep conditions revealed that phosphorus stress strongly advanced QS activation and lowered the biomass required for *phz* induction (Fig. 4D,E). Under P-lim, ROIs typically induced within a few hours (mean induction time ∼5 hr) and did so at relatively low biomass, whereas under P-rep conditions induction was markedly delayed (mean ∼32 hr) and required substantially higher biomass at the time of activation (on average 5-fold greater intensity). Induction times were also much more heterogeneous across ROIs under P-rep conditions. Because this reactor is not well mixed, additional transport and resource gradients almost certainly emerge that affect the observed timing that we do not control for (e.g., oxygen depletion in larger pores). In this more realistic geometry, the contrast between P-lim and P-rep induction time appears amplified relative to previous experiments (Figures 2 and 3).

We next asked whether QS induction under P limitation depends on initial cell density within individual water pockets. To quantify this feature, we used the ratio of cell area to water area at t = 1 hr for each ROI as a metric for initial cell density and related it to the time of *phz* induction for each ROI. Across P-lim experiments, induction time decreased systematically with increasing starting density, following a non-linear, approximately power-law relationship (R^2^ = 0.65, Fig. 4F, Fig. S3). Pearson and Spearman correlations gave similar results (r = -0.771, p = 4.14e-20; ρ = -0.839, p = 1.59e-26). Therefore, most of the variation in induction timing is explained by initial cell density, supporting a view in which phosphorus stress and spatial confinement allow cells concentrated in pore water to accelerate phosphate depletion and AHL accumulation to facilitate QS activation.

Taken together, these data demonstrate that the combination of phosphorus stress and spatial confinement enables QS activation in microenvironments that would otherwise restrict signal buildup. We next asked whether lowered QS concentration thresholds could provide an ecological advantage in a community setting.

### Phosphorus limitation confers cooperative and antagonistic quorum-sensing-driven community dynamics

Under nutrient stress, microbial communities can flip between cooperative information exchange (e.g., quorum signals) and antagonistic interference (e.g., antibiotics), reshaping which partners win or lose.

To determine whether phosphorus limitation broadly enhances AHL production and promotes cross-talk among *Pseudomonas* species, we assembled a panel of rhizosphere-associated strains representing both close relatives of *P. synxantha* and more distantly related pseudomonads. Each strain was grown under N-lim, P-lim, or P-rep conditions, cell-free supernatants were then collected to challenge the *P. synxantha* Δ*phzI*–P*_phzA_* reporter, which cannot synthesize AHLs but fluoresces upon QS-activated *phzA* expression (Fig. 5A). Supernatants from multiple donors activated the reporter, with the strongest and broadest induction under P-lim and P-rep, and little to none under N-lim. As expected, controls with the Δ*phzI* donor did not induce, whereas the wild-type donor triggered a robust response. Notably, across the panel, strains carrying homologous *phzIR* operons consistently induced *phzA* under P-lim, suggesting that phosphorus stress broadly elevates AHL output among *Pseudomonas* species. In contrast, *P. aeruginosa* PA14 failed to induce the reporter, consistent with its production of 3-oxo-acyl-HSLs rather than the 3-hydroxy-acyl-HSL(s) recognized by PhzR in *P. synxantha*.

**Figure 5:**
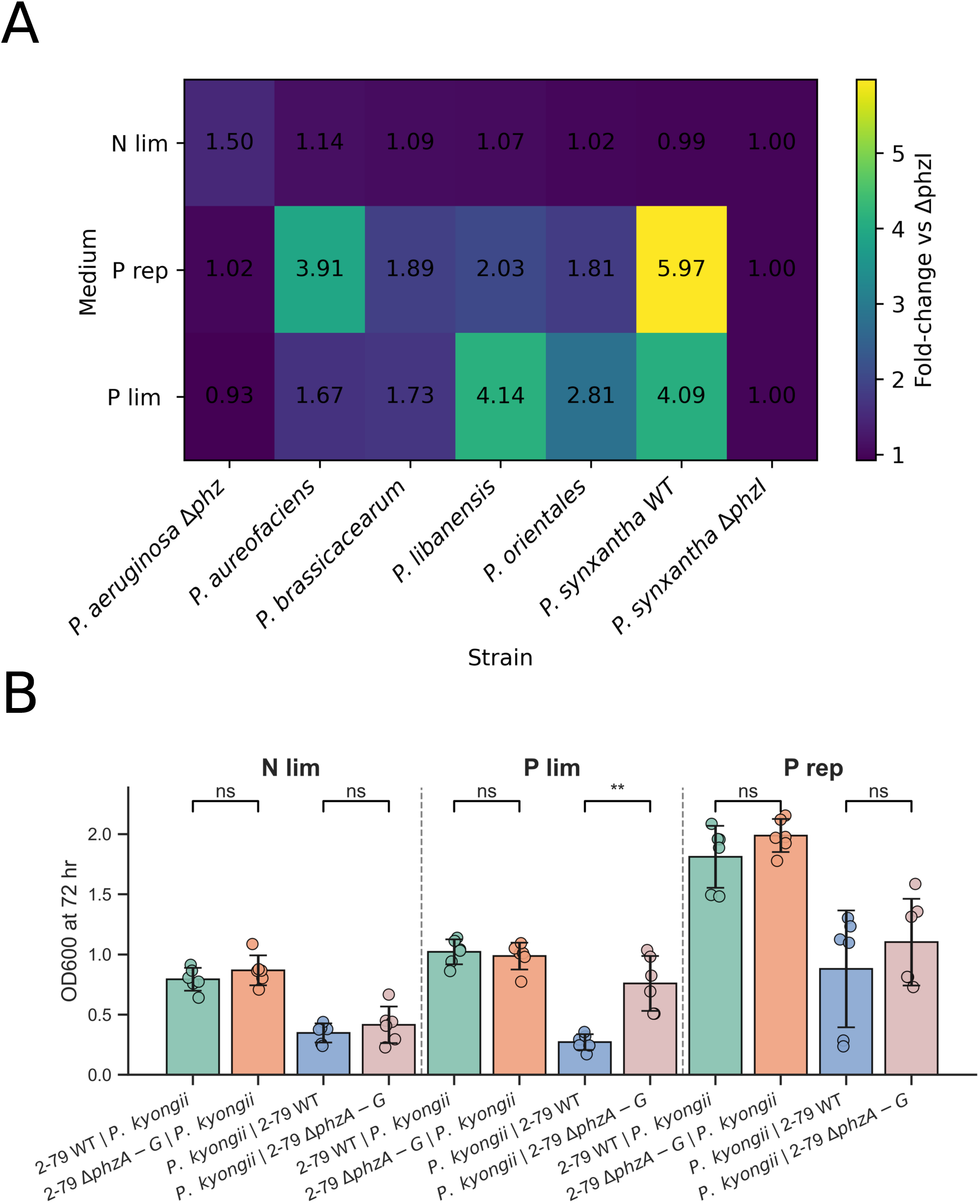
(A) *Pseudomonas* spp. supernatant cross-induction QS assay. Heatmap of fluorescence induction in *P. synxantha* Δ*phzI*-P*_phzA_* across supernatants of selected strains and media conditions at 24 h. Rows represent media (N-lim, P-lim, P-rep); columns are strain supernatants. Each cell shows the fold-change of the chosen metric relative to the reference strain, *P. synxantha* Δ*phzI*, within the same medium. Where shown, numeric labels are fold-change values rounded to two decimals; the *P. synxantha* Δ*phzI* column equals 1 by definition. Fluorescence is normalized by OD600. (mean, *n* = 6) **(B) Competition assays between *P. synxantha* against *Pedobacter kyonggii* under various nutrient regimes**. Bar plots show OD600 at 72 h (mean ± SD, *n* = 6) for four pairings. The first name indicates the strain measured; the second is its pairing partner. Results are shown separately for N-lim, P-lim, P-rep media. *P. synxantha* wild type (2-79 WT) and *P. synxantha* Δ*phzA-G* (2-79 Δ*phzA-G*). Statistical comparisons use Welch’s *t*-test; brackets denote significance (ns, not significant; ** p < 0.01).

Next, we asked whether QS activation under P-lim alters microbial interactions within competitive communities under oxic conditions where phenazines primarily function as antibiotics.^11^ We performed co-culture assays between *P. synxantha* (wild type and Δ*phzA-G*, a phenazine-null mutant) and *Pedobacter kyonggii*, a common rhizosphere isolate. When P-limited, the wild-type strain significantly suppressed *P. kyonggii* growth, whereas the Δ*phzA-G* mutant did not (Fig. 5B). This phenazine-linked advantage was absent under N-lim in the wild-type strain and, interestingly, under P-rep conditions as well. Although the wild type produced PCA under P-replete conditions, induction occurred much later (∼35 h vs ∼16 h in P-lim; see Fig. S5), by which time the co-cultures had already completed most of their growth, leaving little opportunity for inhibition to manifest. Thus, the phenotype likely reflects the timing of phenazine onset relative to culture growth.

These findings indicate that phosphorus limitation broadly elevates AHL signaling across *Pseudomonas* spp., priming QS in *P. synxantha*. Moreover, phenazine production under phosphorus stress suppresses growth of *P. kyonggii*.

## Discussion

Our study reveals that phosphorus limitation rewires the quorum-sensing (QS) circuitry of *Pseudomonas synxantha* 2-79, lowering the effective signal concentration required for autoinduction and enabling phenazine biosynthesis at conditions corresponding to much lower densities (OD ≈ 0.08 in liquid cultures). Phosphorus stress reduces the acyl-homoserine lactone (AHL) concentration required for activation by roughly an order of magnitude under our tested conditions, thereby heightening cellular sensitivity to QS signals. This response depends on the PhoBR two-component system: inactivation of PhoBR abolishes phenazine induction even when AHL levels are high, demonstrating that phosphorus stress provides a necessary regulatory input. In structured habitats, where mass transfer alters QS molecule accumulation, phosphorus limitation emerges as a key modulator of QS activation. In colony biofilms, phenazine induction times under P limitation are strongly affected by agar volumes and biofilm density, implicating biofilm retention and diffusive losses in achieving threshold AHL concentrations.

Exogenous AHL addition collapses this concentration requirement but still requires a physiological state primed for induction (i.e., P limitation or higher cell densities under P replete). Confinement in micropores creates microenvironments that retain signals and enable localized QS activation, again showing that physical structure and nutrient state jointly set QS thresholds. These environmentally-tuned threshold shifts have ecological consequences. Under phosphorus limitation, *P. synxantha* inhibits growth of the rhizosphere isolate *Pedobacter kyonggii* through phenazine production, a benefit lost in the phenazine-null mutant and under nitrogen limitation or phosphorus-replete conditions. Early phenazine synthesis may thus serve dual functions in resource competition and phosphorus acquisition, coordinated through stress-responsive signaling.

Our data argue against “concentration-only” or “density-only” views and instead support an AND-gate model for QS activation: (i) sufficient signal accumulation (production versus diffusion, advection, sorption, and decay) to cross a threshold and (ii) a physiological readiness state that licenses responsiveness. Density remains a useful proxy because per-cell production scales with biomass and high density often coincides with a physiological readiness, but other regulatory programs can shift the effective threshold, explaining low-density induction under nutrient stress. In our system, QS signal responsiveness is tunable by nutrient status, PhoBR under phosphorus limitation, so signal is necessary but not sufficient. This framework reconciles our observations: under P-replete + excess AHL, low-density cultures fail to induce despite ample signal, indicating physiological unreadiness; responsiveness emerges by mid- to late-exponential growth as global regulators (e.g., Gac/Rsm) and stress programs rise. Under P limitation, induction synchronizes with phosphorus stress. Exogenous AHL enables induction at comparatively lower cell densities, with the activation density tunable by the imposed Pi level. Again, though, induction did not occur until P limitation was achieved.

This model aligns with prior work. Overactivation of the stringent response via RelA in *P. aeruginosa* prematurely activates RpoS, LasR, RhlR and autoinducer synthesis, driving LasB production independent of density, i.e., physiological readiness upstream of signal thresholds.^65^ Exogenous PQS (an autoinducer) addition can override cell-density dependence but not growth-phase dependence, underscoring the physiological readiness requirement.^66^ More broadly, nutrient limitation and growth rate jointly modulate extracellular signal accumulation across species, with phosphorus and sulfur limitation disproportionately elevating alkylquinolones even at lower OD.^67^ Our findings extend this signal-accumulation + physiological readiness concept by showing that native PhoBR directly sensitizes an unmodified AHL-based QS network in a rhizosphere pseudomonad, reframing QS as an environmentally tuned concentration-sensing mechanism whose effective activation threshold is set jointly by autoinducer concentration and regulatory state. Related frameworks^43^ reach similar conclusions; nevertheless, proliferating new labels for QS risks semantic confusion that can imply a single dominant driver.^36^ Instead, we favor a generalizable AND-gate view that explains QS outputs in sparse, fragmented soils and highlights practical levers (e.g., nutrient availability, confinement, and mass transfer) for engineering compatible microbial communities and biosensors in heterogeneous environments.

## Methods

### Bacteria

The primary strain was *Pseudomonas synxantha* (formerly *P. fluorescens*) 2-79 WT. Isogenic derivatives included Δ*phzI* (AHL synthase null), Δ*phzA-G* (phenazine-null), and Δ*phoB* (phosphorus-stress response regulator null). Reporters carrying transcriptional fusions included P*_phzA_*–mNeonGreen (QS/phenazine biosynthesis), P*_phoA_*–mNeonGreen (phosphorus stress), P*_phzA_*–GFP, P*_tac_*–mCherry (QS/phenazine biosynthesis, density metric) and P*_PA10403_*-mNeonGreen (density metric). Donor/competitor strains for crosstalk and competition assays included *Pseudomonas aureofaciens* 30-84, *P. brassicacearum* 106, *P. libanensis* 84, *P. orientalis*, *P. aeruginosa* Δ*phz1/2* and *Pedobacter kyonggii*. Sources, identifiers, and genotypes are listed in Table S2.

#### Culture conditions

Unless noted, overnight cultures of *Pseudomonas* strains were propagated at 30 °C, 250 rpm in LB (Miller) for 16-18 h; *E. coli* DH10B/SM10 were grown at 37 °C. Experimental assays used defined MOPS medium (below). Antibiotics were used only for selection (gentamicin 30 µg mL^-1^, tetracycline 15 µg mL^-1^). All experiments with reporters were antibiotic-free. Strains were stored as −80 °C glycerol stocks (20% v/v) and streaked fresh for each experiment.

### Media compositions and nutrient-regime definitions

#### MOPS defined medium

Per liter: 25 mM MOPS (pH 7.0; adjusted with 1 M NaOH), 55.5 mM glucose, 16 mM NH₄Cl, 7 mM KH₂PO₄, 0.41 mM MgSO₄·7H₂O, 0.68 mM CaCl₂·2H₂O, Aquil-based trace metals^68^ with Fe 10 µM and EDTA 100 µM. KH₂PO₄ 7 mM (P-rep/N-lim) or 25–100 µM (P-lim); NH₄Cl 16 mM (P-rep/P-lim) or 2 mM (N-lim). No-N and No-N/P buffers omitted NH₄Cl and NH₄Cl/KH₂PO₄, respectively. MOPS agar consisted of MOPS medium with 1.0% (w/v) noble agar.

#### Plant medium

0.5× MS (Sigma-Aldrich M5519), 0.6% (w/v) Phytagel (Sigma-Aldrich P8169), pH 5.8 (KOH); P-lim omits KH₂PO₄.

#### Phosphate-sorbing (P-HFO) condition

Hydrous ferric oxide (HFO) was synthesized, phosphate was then sorbed and sequestered (P-HFO). P-HFO was added to phosphorus deficient plant medium (see below) at 1.6 g/L. See Note S1 for P-HFO preparation.

#### MES defined medium

Per L: 50 mM MES pH 5.8 (KOH), 50 mM sucrose, salts as above. KH₂PO₄ 7 mM (P-rep/N-lim) or 100 µM (P-lim); NH₄Cl 16 mM (P-rep/P-lim) or 2 mM (N-lim).

### Reporter and mutant generation and verification

Transcriptional fusions were constructed using the mini-Tn7 system.^69^ Promoter (P*_phzA_*) was cloned upstream of mNeonGreen in pJM220 by Gibson assembly and transformed into *E. coli* DH10B. Gentamicin-resistant transformants (30 µg mL^-1^) were verified by colony PCR and Sanger sequencing. Mini-Tn7 elements and pTNS1 helper (TnsABCD; delivered from *E. coli* SM10/pTNS1) were introduced into electrocompetent *P. synxantha* 2-79. Transconjugants were selected on LB-gentamicin and screened by junction PCR for single-copy insertion at attTn7 (downstream of *glmS*). Reporter function was validated in plate-reader assays (P*_phzA_* induction under QS-permissive conditions). Plasmids and primers are listed in Tables S3–S4.

In-frame deletion mutants were constructed using a suicide plasmid. Approximately 1-kb regions upstream and downstream of *phzA-G* and *phzI* were PCR-amplified and assembled with SmaI-digested pEX18Gm by Gibson assembly. The plasmid was electroporated into *E. coli* DH10B, and transformants were selected on LB agar with gentamicin (30 µg mL^-1^). Correct constructs were verified by PCR and introduced into *P. synxantha 2-79* wild-type strains via triparental conjugation. Merodiploids were selected on Vogel–Bonner minimal medium (VBMM) containing gentamicin (30 µg mL^-1^), followed by counter-selection on sucrose medium (5 g L^-1^yeast extract, 10 g L^-1^tryptone, 100 g L^-1^sucrose). Colonies in which homologous recombination was successful were identified by PCR. Deletions were confirmed by diagnostic PCR across the targeted loci and by phenotype: Δ*phzA-G* lacked ABS367nm (PCA readout) signal and lacked phenazine product, verified via LC-MS analysis. Δ*phzI* required exogenous AHL for induction and lacked AHL production, verified via LC-MS/MS analysis.

### Plant-microbe reporter induction assay

*Brachypodium distachyon* Bd21-3 seeds were dehusked, sterilized with chlorine gas (see Note S2), plated on 0.6% Phytagel with 0.5× MS, and vernalized (4 °C, 4 d). Seedlings were grown 2 d (16 h light/8 h dark, 25 °C), then immersed 30 min in 0.5× MS lacking phosphate (P-lim) containing *P. synxantha* 2-79 at OD_600_ = 0.1. Seedlings were transferred into 1 cc syringes filled with 0.6% Phytagel prepared with 0.5× MS under the indicated phosphorus condition (P-lim, P-rep, or P-HFO) and incubated vertically for 2 d prior to imaging.

Seedlings embedded in phytagel were mounted so that roots+phytagel were submerged in sterile 0.5× MS buffer. Imaging used a Zeiss Lightsheet Z.1 with EC Plan-Neofluar 5×/0.16 air objective. Dual-channel acquisition captured GFP/mNeonGreen (488 nm ex, 498 nm em) and mCherry (561 nm ex, 571 nm em) on separate cameras with a dual-band emission filter (Zeiss LBF 488/561). Z-stacks were collected at 5 µm step, XY pixel 0.93 µm, 100 ms exposure per slice; laser power was held constant across conditions (20% at 488 and 561 nm). Maximum intensity projections (MIPs) were generated for analysis.

All analysis used Fiji/ImageJ with a custom macro. Red channel (P*_tac_*–mCherry) was median-filtered (r = 4), background-subtracted (rolling ball 50 px), converted to 8-bit, and locally thresholded (Bernsen, r = 15) to build masks. Objects were defined by particle analysis (min size 20 px) and applied to original red/green images to extract integrated density (IntDen) and area. Mean intensity per object was computed as IntDen/area for both channels. CSVs were exported for Python analysis.

### Planktonic 96-well kinetic experiments

Overnight cultures of *P. synxantha* 2-79 (WT, Δ*phzA-G*, Δ*phzI*, Δ*phoB,* Δ*phzI–*P*_phzA_*) were washed 3× in MOPS containing 10 µM phosphate, and inoculated into 96-well plates at OD_600_ = 0.01. Plates were incubated in a Tecan Spark 10M (30°C; orbital shaking amplitude 2.5 mm, 216 rpm) with reads every 15 min for 24 h. OD_600_ was read at 600 nm; ABS367 was read at 367nm for PCA-associated absorbance; mNeonGreen at Ex/Em 485/535 nm. Induction time was approximated from fluorescence time series by manual curation. ODs at induction were obtained by linear interpolation of OD_600_ (t) at that time.

For the AHL-supplemented dose-response assays, 3-OH-C6-HSL (Adipogen; ≥98%) stock was prepared with methanol (final solvent ≤0.02% v/v; vehicle controls showed no effect from solvent carryover). Dose ranges spanned sub-nM to tens of µM (specified per figure).

For the no-growth supplementation assay, Δ*phzI* overnight culture was washed (3×) in the respective treatment buffer (No-N or No-N/P) were inoculated at OD_600_ = 0.1. For the growth-permissive supplementation assays, Δ*phzI* overnight culture was washed (3×) in MOPS containing 10 µM phosphate and inoculated at OD_600_ = 0.01. Responses were baseline-corrected to 0 µM within condition and fit with a two-parameter Hill (log-logistic) model (top = 1, bottom = 0) to estimate EC_50_ with 95% CIs.

### AHL quantification by LC–MS/MS

Overnight cultures of *P. synxantha* 2-79 (WT, Δ*phzI*) were washed 3× in MOPS, then grown in glass culture tubes (30 °C, 250 rpm) under P-rep, P-lim, and N-lim conditions for 22 h. Cultures were then pelleted, supernatants were collected, filtered through a 0.22 µm PES filter, diluted 1:1 in methanol, incubated for 1 h at -20°C, pelleted, then supernatants were collected before analyzing.

Analytical standards (3-OH-C6-HSL, 3-OH-C8-HSL, 3-OH-C10-HSL, and 3-oxo-C8-HSL; ≥95–98% purity; Adipogen/Cayman) were prepared by solubilizing powders in methanol as 1-10 g L^-1^ primary frozen stocks (−20 °C). A 100 µM all analyte mix was made in methanol with 0.1% formic acid. Calibration standards were prepared by diluting the analyte mix in methanol with 0.1% formic acid.

A Waters (Milford, MA) Acquity Premier UPLC was fitted with a HSS T3 column and coupled to a Xevo Q-TOF G2S. Mobile phase A was 0.1% formic acid in water and mobile phase B was 100% acetonitrile. The initial mobile phase composition was 2.5 % B for 0.5 minutes with a linear gradient to 95% B ending at 5 minutes. After a 0.5 minute hold (5.5 min) the composition was returned rapidly to the initial conditions (5.6 min) and held for an additional 0.25 minutes (to 5.74 minutes). The gradient was cycled without interruption during any acquisition. The flow rate was held constant throughout at 0.3 mL/min, the samples held at 10 °C, the column temperature maintained at 40 °C, and the injection volume was 4 µL. The 2.1 mm × 50 mm × 1.8 µm column was used without a guard column.

The ion source block was held at 120 °C, capillary voltage was 3 kV, desolvation gas flow and temperature were 600 lph and 500 °C, the cone gas flow as 15 lph, and the source offset was 80 V. Separate mass spectral data sets were collected in both MS mode from 50-1200 m/z and using scheduled MS2 for parallel reaction monitoring of the individual targeted analytes. MS data was acquired in positive sensitivity mode (approximately 18,000 resolution). The mass axis was calibrated against sodium formate and was confirmed daily. Leucine enkephalin was used as lock mass reference during acquisition.

Targeted analysis was performed using the Quanlynx program within the Masslynx software suite used to operate the instrument.

### Sessile early-stage colony biofilm experiments

Overnight cultures of *P. synxantha* 2-79 (WT, P*_phoA_*, P*_phzA_*, P*_PA10403_*) were washed 3× in MOPS containing 7 mM phosphate and adjusted to OD_600_= 0.05. A 5 µL droplet was spotted onto agar plates, air-dried 1 h, and placed in a custom humidity-controlled platform. The agar plates contained either 2 mL or 5 mL of agar. For P-lim plates, phosphorus was omitted from the MOPS agar; the only source of phosphorus was the 5 µL inoculation drop that contained 7 mM phosphate. P-rep consisted of MOPS agar at 7 mM. Images were acquired hourly for 36 h.

Time-lapse imaging of bacteria was performed using a previously reported complex-field and fluorescence microscopy system based on the aperture scanning technique (CFAST).^48^ Samples were mounted in a 12-well plate and positioned on a motorized XY scanning stage (Thorlabs MLS203-1) to enable automated, parallel scanning of multiple wells.

To minimize evaporation of the growth medium during the two-day imaging experiments, the 12-well plate was placed in a custom 3D-printed holder containing wet paper towels to maintain high local humidity. To maintain focus while minimizing photobleaching and phototoxicity, 0.5-μm red fluorescent beads (580/605 nm, ThermoFisher F8812) were pre-mixed with the bacterial suspension at a 1:10⁵ dilution from stock. These beads served as markers for focal plane tracking. The beads were excited with a 568 nm laser (Coherent Sapphire 568 LP), and the emitted fluorescence was filtered using a dichroic mirror (Semrock Di01-R488/561) and a bandpass filter (Semrock FF01-R523/610).

A 488 nm laser (Thorlabs LP488-SF20G) was used both for complex-field illumination and for excitation of mNeonGreen fluorescence. The complex field data comprising both amplitude and quantitative phase was used to quantify the area coverage of bacterial colonies. Fluorescence intensity was extracted from the mNeonGreen images and subsequently normalized using brightfield images to compensate for variations in excitation power due to differences in medium thickness across the sample.

### Microfluidic soil-analog reactor experiments

Devices followed a silicon-etched porous design^64^ and fabrication process (see Note S3). Overnight culture of *P. synxantha* 2-79 (dual reporter, P*_phzA_*-P*_tac_*) was washed 3× in MOPS P-lim (100 µM) or P-rep medium and adjusted to OD_600_= 0.05 for inoculation into the reactor. Cells were inoculated at a flow rate of 0.1 mL min^-1^ for 3 minutes. Air was then injected and generated heterogeneous water saturation (connected high-saturation domains vs confined pockets). After initial displacement, air was continuously injected at a flow rate of 0.01 mL min^-1^. Imaging was conducted for 16-40 h depending on experiment.

All imaging was performed on a Nikon Ti2 widefield epifluorescence microscope. Time-lapse series used a Plan Apo 20×/0.75 dry objective. GFP was excited with a 488-nm LED and collected with a FITC filter set; mCherry was excited with a 561-nm LED and collected with a TRITC filter set. Images were collected every hour from multiple stage positions.

Custom MATLAB code was used to segment bacterial cells based on the red channel (constitutive reporter) and pockets of air based on the green channel (induction of phenazine production). To identify cells, background subtraction was applied to the original image followed by thresholding using a fixed intensity value. Objects smaller than 5px were discarded. To segment pockets of air, we first identified and removed signal from induction, which would otherwise interfere with the detection. For this we segmented fluorescing cells by simple thresholding, expanded their area by dilation (10px) and then replaced intensity values within this area with the background intensity. Following this, air pockets were segmented by smoothing and background subtraction followed by thresholding. Areas smaller than 1000px were discarded and holes within air pockets smaller than 50000px were closed. When necessary, segmentation of air pockets was corrected manually. Intensity ratios were based on background-corrected intensities. Induction time vs cell density was fit to a power-law model using log–log linear regression; goodness-of-fit (R²) and CI on parameters are reported.

### Supernatant cross-induction assay

Overnight cultures of *Pseudomonas* strains were washed 3× in MOPS (P-lim and N-lim) medium and inoculated at OD_600_ =0.01 into glass culture tubes (30 °C, 250 rpm) under P-rep, P-lim, and N-lim conditions for 22 h. Cultures were then pelleted, and supernatants were collected and filtered through a 0.22 µm PES filter. The cell-free supernatant was then mixed 1:1 with appropriate medium (i.e., P-rep, P-lim, N-lim) for cross-induction assay.

An overnight culture of *P. synxantha* 2-79 Δ*phzI-*P*_phzA_* was washed 3× in MOPS (P-lim and N-lim) and inoculated at OD_600_ =0.01. A 96-well plate assay was then conducted with the cell-free supernatant mixture following the “Planktonic 96-well kinetic experiments” parameters. Fold-change for RFU/OD_600_ relative to the reference strain *P. synxantha* Δ*phzI* within each medium was then computed. The resulting scores were plotted as a single heatmap.

### Pairwise competition assays

Co-culture growth and induction dynamics were monitored using a two-chamber co-culture system that allows fluidic exchange while physically separating populations (Duet Co-Culture System, Cerillo). *P. synxantha* (WT or Δ*phzA-G*) and *P. kyonggii* overnight cultures were washed 3× in MES (P-lim and N-lim) medium. Cells were then loaded to either side of the chamber at a starting OD_600_ of 0.01 under N-lim, P-lim, or P-rep media. Duets have the footprint of a 96-well plate, thus the assay was conducted following “Planktonic 96-well kinetic experiments” instrument parameters. Phenazine-dependent suppression was assessed by comparing WT vs Δ*phzA-G*. Statistics used Welch’s t-test with two-sided α = 0.05. Note that competition assays were run in MES-defined medium (pH 5.8) with sucrose as the carbon source. Sucrose was chosen because *P. synxantha* 2-79 does not utilize it; on glucose, *P. synxantha* rapidly overgrows *Pedobacter kyonggii*, obscuring interaction effects. Using sucrose maintains competitive viability of *P. kyonggii* and shifts *P. synxantha* toward relying on metabolites released by P*. kyonggii*, enabling resolution of phenazine-dependent outcomes. The pH was set to 5.8 (buffered with MES) to approximate mildly acidic rhizosphere conditions and because PCA is more membrane-permeant—and thus more potent—at lower pH.

## Acknowledgements

We acknowledge the Resnick Ecology and Biosphere Engineering Facility and Caltech Biological Imaging Center for the training and use of confocal and light sheet fluorescence microscopes. We are grateful to the Resnick Sustainability Institute and National Institute of Health (2R01AI127850-06A1) for enabling resources and financial support. R.E.A. was supported by an NSF Postdoctoral Research Fellowship in Biology (Grant No. 2209379). H.J. was supported by a Helen Hay Whitney Postdoctoral Research Fellowship.

## Author contributions

R.E.A., and D.K.N. designed research; R.E.A., H.J., O.Z., R.L.A., and N.D. performed research; R.E.A., H.J., O.Z., R.L.A., and N.D. analyzed data; and R.E.A., H.J., O.Z., R.L.A., N.D., D.M., C.Y., and D.K.N. wrote the paper.

