## Supplemental Tables and Figures for "Phosphorus stress and spatial confinement lower the threshold for quorum-sensing activation of redox-active metabolite production in *Pseudomonas synxantha*"

**Supplemental Material**

**Table S1: Summary statistic for LSFM imaging**

| Condition | Channel | Count | Mean | Std | SEM | Min | Max | IQR | Skew | Kurtosis |
|---|---|---|---|---|---|---|---|---|---|---|
| dual P-hfo | Green | 469 | 0.13844 | 0.14716 | 0.0068 | 0.0075 | 1 | 0.13163 | 2.58119 | 8.65501 |
| | Red | 469 | 0.11688 | 0.12996 | 0.006 | 0.00044 | 0.87738 | 0.12079 | 2.45011 | 7.89059 |
| dual P-lim | Green | 233 | 0.06059 | 0.07714 | 0.00505 | 0.01874 | 0.77512 | 0.01942 | 4.92864 | 34.71869 |
| | Red | 233 | 0.05101 | 0.10307 | 0.00675 | 0 | 0.83051 | 0.04177 | 5.23136 | 30.49959 |
| dual P-rep | Green | 982 | 0.01605 | 0.01017 | 0.00032 | 0 | 0.13305 | 0.0098 | 3.39508 | 25.84241 |
| | Red | 982 | 0.1153 | 0.13696 | 0.00437 | 0.00018 | 1 | 0.09841 | 2.89134 | 10.31037 |

**Table S2: List of strains**

| Species/Strain | Genotype / Description | Purpose | Source / Reference |
|---|---|---|---|
| *Pseudomonas synxantha 2-79* | Wild type | Baseline physiology; AHL & PCA production | Gift of D. Mavrodi |
| *Pseudomonas synxantha 2-79 ΔphzI* | AHL synthase knockout | AHL complementation; dose–response; cross-induction reporter host | This study |
| *Pseudomonas synxantha 2-79 ΔphzA-G* | Phenazine-null (operon deletion) | ABS367 baseline; phenazine-dependent phenotypes | This study |
| *Pseudomonas synxantha 2-79 ΔphoB* | PhoBR regulon (*phoB*) deletion | QS induction under P-lim; PhoRB requirement | Lab collection[1] |
| *Pseudomonas synxantha* 2-79 attTn7::$P_{\mathit{phzA}}$–mNeonGreen | Single-copy chromosomal reporter | QS/phenazine transcription reporter | Zhang, Alcalde et al.[2] |
| *Pseudomonas synxantha 2-79* attTn7::$P_{\mathit{phoA}}$–mNeonGreen | Single-copy chromosomal reporter | Phosphate stress reporter | Zhang, Alcalde et al.[2] |
| *Pseudomonas synxantha 2-79* attTn7::$P_{\mathit{PA10403}}$–mNeonGreen | Single-copy chromosomal reporter | Constitutive reporter used as cell density proxy in colony biofilm assays | Zhang, Alcalde et al.[2] |
| *Pseudomonas synxantha 2-79* attTn7::$P_{\mathit{tac}}$–mCherry pPROBE:: $P_{\mathit{phzA}}$-GFP | Single-copy chromosomal and multi-copy plasmid reporter | Dual reporter as density proxy and *phz* induction. Imaging masks; phenazine transcription reporter | Gift of D. Mavrodi |
| *Pseudomonas synxantha 2-79* Δ*phzI*, attTn7::$P_{\mathit{phzA}}$–mNeonGreen | AHL-blind transcription reporter | Dose-response curves, Supernatant cross-induction assay | This study |
| *Pseudomonas aeruginosa PA14 Δphz1/2* | Phenazine-null (both phz operons) | Donor for AHL cross-induction | Lab collection[3] |
| *Pseudomonas aeruginosa PA14* | Wild type | Donor for AHL cross-induction | Lab collection |
| *Pseudomonas aureofaciens 30-84* | Rhizosphere isolate | Donor for AHL cross-induction | Lab collection |
| *Pseudomonas brassicacearum 106* | Rhizosphere isolate | Donor for AHL cross-induction | Gift of D. Mavrodi |
| *Pseudomonas libanensis 84* | Rhizosphere isolate | Donor for AHL cross-induction | Gift of D. Mavrodi |
| *Pseudomonas orientalis* | Rhizosphere isolate | Donor for AHL cross-induction | Lab collection[4] |
| *Pedobacter kyonggii* | Rhizosphere isolate | Competition assays | Lab collection[4] |
| *Escherichia coli* DH10B | Cloning host | Assembly of mini-Tn7 constructs | Lab collection |
| *Escherichia coli* SM10 (pTNS1) | Helper for TnsABCD transposition | transposon for mini-Tn7 delivery | Lab collection[5] |
| *Escherichia coli* HB101 (pRK2013) | Helper with pRK2013 | for mobilization of non-self-transmissible plasmids. | Lab collection[5] |

**Table S3: List of plasmids**

| Plasmid | Purpose | Source |
|---|---|---|
| pJM220 | Single-copy chromosomal insertion at attTn7 | Lab collectoin[6] |
| pTNS1 | TnsABCD transposon for mini-Tn7 delivery | Lab collection[5] |
| pRK2013 | Mobilization of non-self-transmissible plasmids. | Lab collection[5] |
| pEX18Gm | allelic-exchange (suicide) vector | Gift of D. Mavrodi |
| pEX18Gm [Δ*phzI*] | Generate Δ*phzI* | This study |
| pEX18Gm [Δ*phzA-G*] | Generate Δ*phzA-G* | This study |
| pJM220::$P_{phzA}$–mNeonGreen | Inducible promoter driving mNeonGreen expression | Zhang, Alcalde et al.[2] |
| pJM220::$P_{PA10403}$–mNeonGreen | Constitutive promoter driving mNeonGreen expression | This study |

**Table S4: List of primers**

| Sequence (5′→3′) | overhang | Amplicon |
|---|---|---|
| gcttgcatgcctgcaggtcgactctagaggatccccTGATCACGCCAATTTCCTCC | pMQ30 BB | upstream *phzA-G* operon fwd |
| AGGCAGTGTTCTCCTTAGTTG | - | upstream *phzA-G* operon rev |
| gcgcgtaatttcagtcaactaaggagaacactgcctATAGCCGTGACCCATCTCG | upstream *phzA-G* operon | downstream *phzA-G* operon fwd |
| gctatgaccatgattacgaattcgagctcggtacccCACACCCAGCAGTTGTCG | pMQ30 BB | downstream *phzA-G* operon rev |
| agctatgacatgattacgaattcgagctcggtacccGGCCAGCAGCCCAGGCAC | pMQ30 BB | upstream *phzI* operon fwd |
| GGCAGGGGATTCCTTGGGGGAG | - | Upstream *phzI* operon rev |
| accgcacttcccccctcccccaaggaatcccctgccTAACGGCCGAGTCGTCCTCTTG | upstream *phzI* operon | downstream *phzI* operon fwd |
| gcttgcatgcctgcaggtcgactctagaggatccccGTAGTGGCCCAGGGCGGG | pMQ30 BB | downstream *phzI* operon rev |

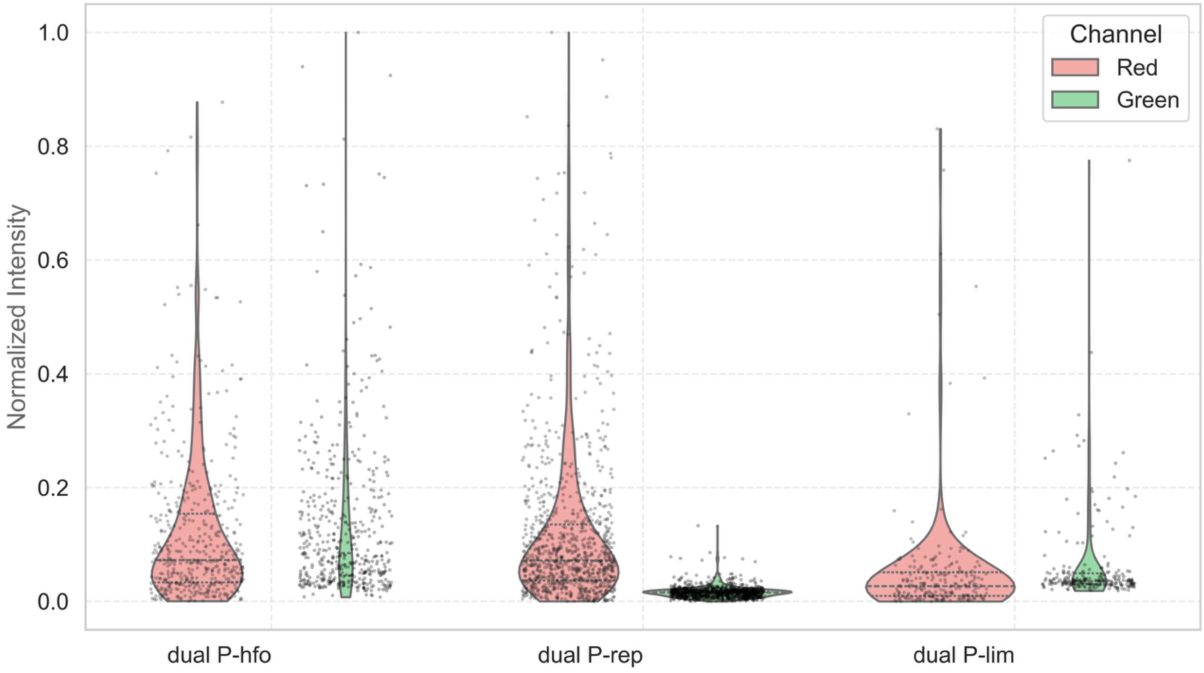

**Figure S1: Violin plots with overlaid strip plots showing normalized fluorescence distributions for each ROI across phosphorus conditions.** Fluorescence was measured in both red (constitutive; $P_{tac}$) and green (phenazine biosynthesis; $P_{phzA}$) channels. Violin plots illustrate the full distribution of normalized ROI-level fluorescence intensities across four conditions: P-HFO, P-rep, and P-lim. Overlaid strip plots reveal individual ROI values, allowing visualization of rare, high-fluorescence outliers that contribute to the observed skew and kurtosis in distribution. These plots emphasize that strong phenazine reporter activation occurs only in P-lim and P-HFO conditions, often in sparse but highly induced subpopulations.

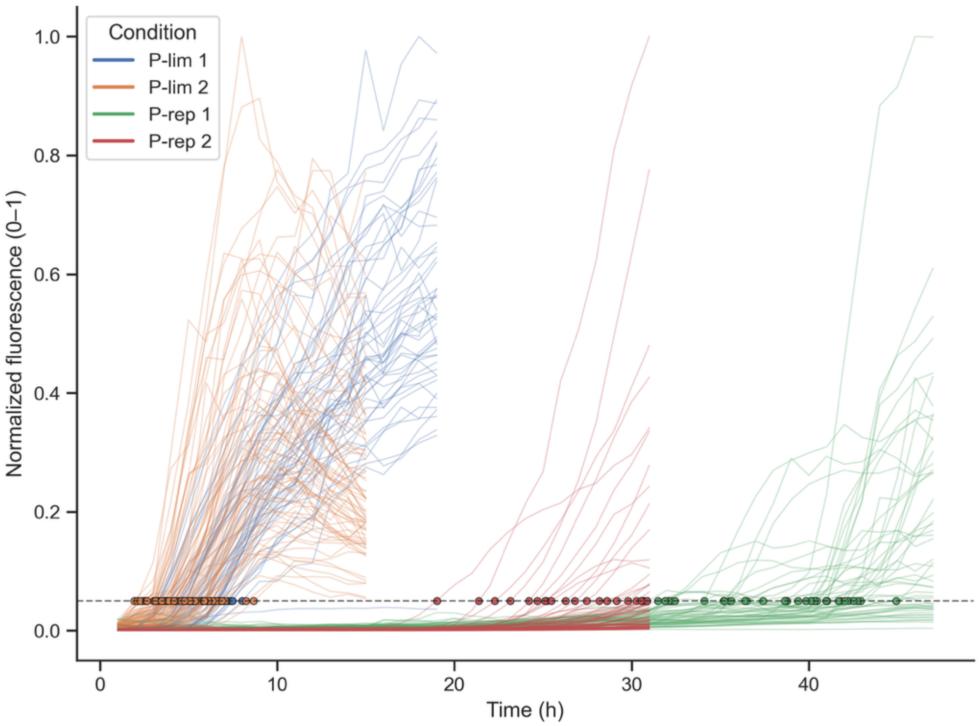

**Figure S2: Time-resolved reporter induction dynamics across phosphorus treatments.** Normalized green-channel fluorescence (integrated fluorescence per ROI min–max scaled to 0–1 within each experiment) is plotted over time for individual ROIs in P-lim 1–2 and P-rep 1–2 microfluidic reactors. Each thin line represents a single ROI trajectory, colored by condition. The horizontal dashed line marks the induction threshold (normalized fluorescence = 0.05), and filled circles indicate the estimated induction time for trajectories that cross this threshold. These induction times are used for the summary comparisons in subsequent panels.

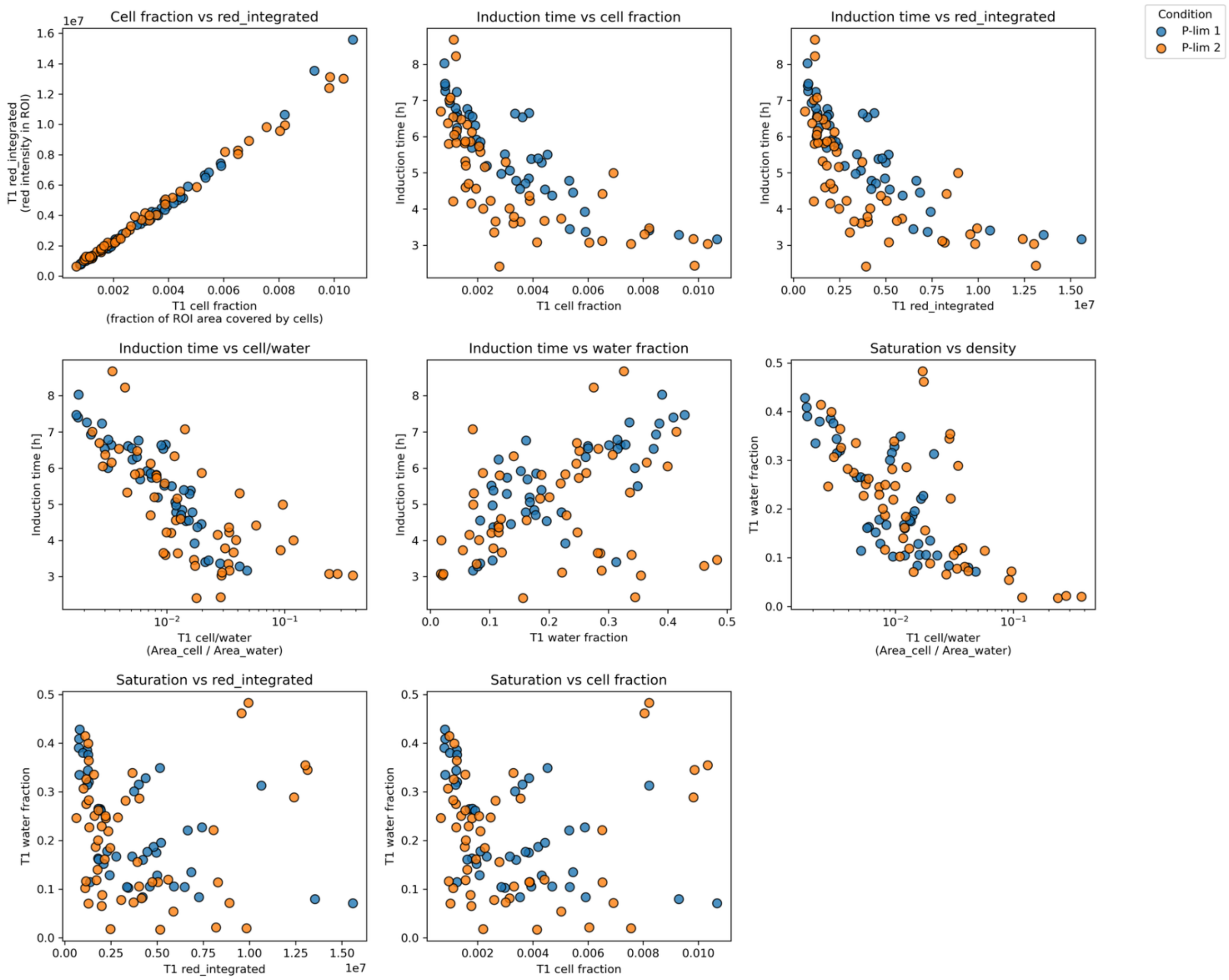

**Figure S3: Relationships between early pore-scale structure and induction time under phosphorus limitation.** Scatter plots show pairwise relationships between t= 1hr (T1) metrics (i.e., cell fraction (area of cells per area ROI), red-integrated (red intensity per ROI) ), water fraction (area of water per area ROI) and cell-to-water area ratio (cell fraction/water fraction) and reporter induction time for P-lim 1 (blue) and P-lim 2 (orange) microfluidic experiments. Each point corresponds to a single ROI. Axes using the cell/water metric are plotted on a logarithmic x-scale to highlight variation in local cell density in pore water. Together, the panel compares how initial cell coverage, red fluorescence, pore-water cell density, and water saturation relate to one another and to the timing of induction across replicate phosphorus-limited reactors.

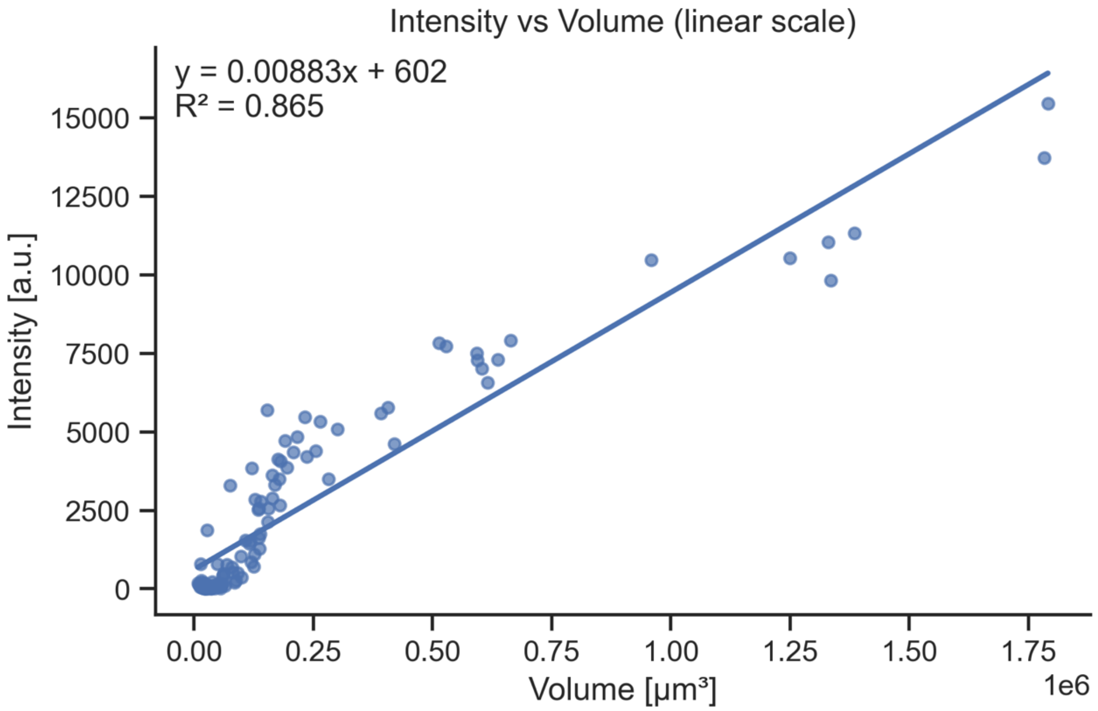

**Figure S4: Intensity scales with object volume.** (A) Linear-scale scatter of per-object fluorescence intensity (a.u.) versus object volume (μm³) with ordinary least-squares best-fit line; inset shows fit equation and $R^2$. Each point is a single segmented region of interest. Volume is the reconstructed 3D fluorescence volume obtained from CFAST's 3D fluorescence capability. Intensity is the wide field intensity value obtained from the same readout. 3D fluorescent volume measurements cannot be captured after cells completely cover the region of interest (ROI), therefore we rely on the intensity measurement for data in Figure 3D.

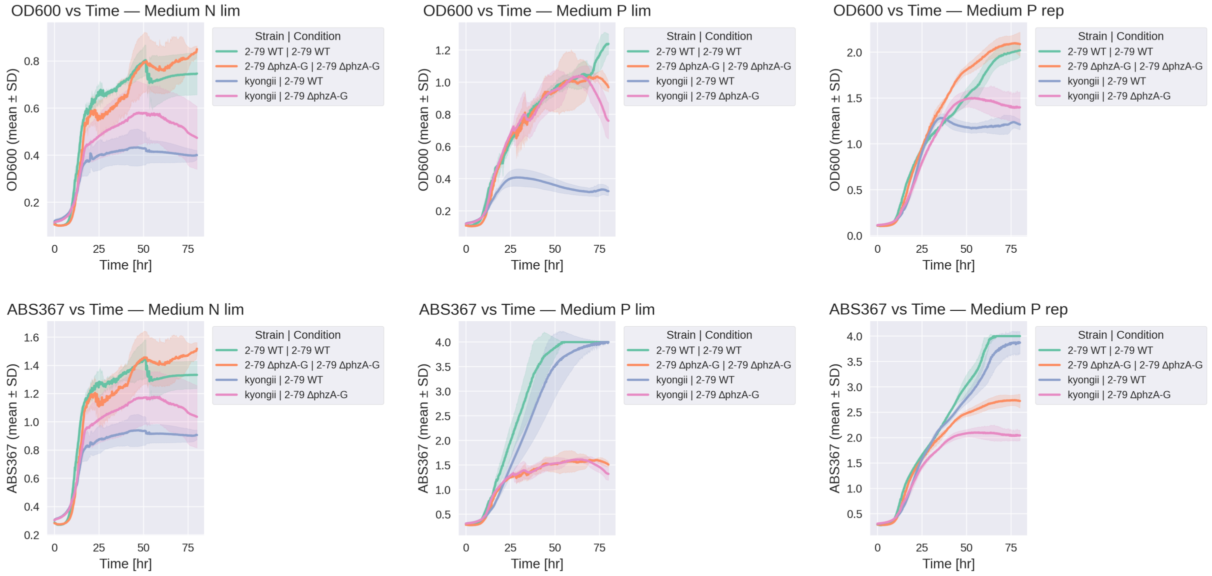

**Figure S5: Growth and phenazine-associated absorbance across nutrient regimes.**
Six-panel composite showing OD600 (top row) and ABS367 (bottom row) versus time for N-limited (N-lim), P-limited (P-lim), and P-replete (P-rep) media. Curves are colored by Strain | Condition, with solid lines indicating the mean across wells and shaded ribbons indicating ± SD. ABS367 serves as a proxy for PCA (phenazine-1-carboxylic acid) production. Time windows reflect the plotted range used in analysis; tick labels denote hours since inoculation. Note that under P-lim conditions, ABS367 rises earlier and at lower densities than under P-rep, consistent with phosphate-stress-sensitized QS activation observed elsewhere in the study. Legends list groups as “Strain | Condition” (e.g., *2-79 WT | 2-79 WT*, *kyonggii | 2-79 ΔphzA-G*). One of two biological replicates represented in main text, n=3.

**Note S1: Hydrous ferric oxide preparation**

Hydrous ferric oxide (HFO) was synthesized following an established protocol.[7] HFO was suspended at 1.6 g $L^{-1}$ in sterile 10 mM KCl and adjusted to pH 5.8 with sterile 1 M HCl, then $KH_2PO_4$ was added to 10 mM final to load phosphate. The suspension was equilibrated for 48 h at room temperature under gentle stirring, maintaining pH 5.8 by checking and correcting at ~0.5, 16, 24, and 48 h. The HFO-P was then pelleted (15,000 × g, 10 min, 4 °C), resuspended in sterile 10 mM KCl (pH 5.8) to the original volume, and this wash was repeated once. A final pellet was resuspended in 0.5× MS phytagel without phosphate (pH 5.8) to 1.6 g $L^{-1}$, and the working suspension was maintained over a stir plate to ensure homogeneity during handling. Experiment was conducted immediately after.

**Note S2: Chlorine gas sterilization of seeds**

Chlorine gas sterilization of seeds was performed in a fume hood with appropriate PPE (gloves, lab coat, eye protection) following previous protocols.[8] Seed tubes (caps open) were placed inside a 7.1 L sealable container along with a 250 mL beaker. To generate chlorine gas, 100 mL household bleach (Clorox; 8.25% sodium hypochlorite) was added to the beaker, followed by 3 mL concentrated HCl (12.1 N); the beaker volume exceeded twice the liquid volume to minimize splashing. The container was sealed immediately, and seeds were exposed to chlorine gas for 1 h. After 1 h, the container was slightly cracked at one corner to vent gas for 3 h, then seed tubes were capped and stored in a cool, dry place.

**Note S3: Microfluidic reactors**

Reactors were microfabricated on silicon by standard photolithography and ICP-RIE to produce a porous landscape of cylindrical pillars (200 µm diameter) with 10 µm depth.[9,10] Inlet/outlet ports (~1 mm diameter) were ultrasonically drilled, and the etched wafer was sealed by anodic bonding to Borofloat-33 glass (etch side up; 900 V, 400 °C). PEEK NanoPort® assemblies (IDEX Corporation) were affixed to the drilled ports with LOCTITE® HP-60 epoxy and coupled to 0.005″ ID PEEK tubing; PEEK/ETFE fittings connected the lines to 5 mL gas-tight syringes (Hamilton 1000 series) driven by a microfluidic syringe pump (Cole-Parmer). Prior to experiments, reactors were cleaned, sterilized, and conditioned by sequential infusion of: Milli-Q $H_2O$ (×2), 1% (w/v) Alconox, Milli-Q $H_2O$, SC-1 (1:1:5 $H_2O_2$(30% w/w):$NH_4OH$(30% w/w):$H_2O$) preheated to 80 °C, Milli-Q $H_2O$, 70% ethanol, autoclaved Milli-Q $H_2O$, and finally MOPS medium.
